# Motor prediction reduces beta-band power and enhances cerebellar-somatosensory connectivity before self-touch to enable its attenuation

**DOI:** 10.1101/2025.07.29.667127

**Authors:** Xavier Job, Lau Møller Andersen, Mikkel C. Vinding, Noa Cemeljic, Daniel Lundqvist, Konstantina Kilteni

## Abstract

Motor control theories suggest that the brain uses forward models to predict self-generated tactile input during voluntary movements, thereby reducing the intensity of reafferent tactile sensations. When one’s own body is the target, this phenomenon is called self-touch attenuation. Although self-touch attenuation is well-documented, it remains unclear how prediction-related neural mechanisms drive attenuation before the self-touch input. We used magnetoencephalography (MEG) to examine the neural correlates of self-touch prediction. 24 human participants (12 female, 12 male) performed a self-touch, and two control tasks. In one control, they received externally generated touch without movement. In the other, the touch was triggered by the participant’s movement, but the hands were spatially misaligned. This manipulation is known to weaken attenuation despite identical tactile input, movement, and task demands, because the sensorimotor context reduces prediction of touch at that body site. Self-touch evoked weaker somatosensory activity (M50 component) than both control conditions. A psychophysics task mirrored the pattern of neural attenuation, as the perception of self-touch was attenuated compared to the two control conditions. To isolate predictive neural mechanisms from general movement-related activity, we subtracted activity from corresponding stimulus-absent trials. Comparing self-touch with misaligned touch allowed us to refine the signal specific to predictive processing in self-touch and revealed greater pre-stimulus beta-band desynchronization and increased cerebellar-to-somatosensory connectivity before self-touch compared to misaligned touch. Our results provide the first evidence of predictive neural activity that shapes the sensory consequences of self-touch, offering insights into the mechanisms through which predictive models modulate somatosensory processing.

**Significance statement:** The brain is thought to predict and attenuate the sensory consequences of self-generated actions, but neural evidence for prediction before sensation has been limited. Using magnetoencephalography, we show that self-touch attenuation is preceded by beta-band desynchronization and increased directed connectivity from the cerebellum to the primary somatosensory cortex. These effects cannot be attributed to movement, as they were reduced in a control condition with similar motor output but lower congruence between the action and tactile consequence, suggesting spatially specific predictive processing. Our study provides the first neural evidence of cerebellar influence on cortical sensory areas before self-touch. These pre-stimulus effects support forward models of sensorimotor control and shed new light on how the brain anticipates and modulates upcoming sensory input.

## Introduction

For centuries, scientists have wondered why we cannot tickle ourselves (Kilteni, 2025). Motor control theories propose that efference copies of motor commands are used by internal forward models, likely involving the cerebellum, to predict the sensory consequences of actions (Blakemore et al., 2000; Kilteni & Ehrsson, 2020; Miall & Wolpert, 1996). These predictions attenuate expected sensory feedback, so self-generated touch applied to the body (self-touch) feels less intense than identical external touch (Bays & Wolpert, 2008; Kilteni, 2023)

Behavioural studies have repeatedly demonstrated self-touch attenuation (Bays et al., 2005, 2006; Blakemore et al., 1999a; Kilteni et al., 2018, 2019, 2021; Kilteni & Ehrsson, 2017, 2020, 2022). Neuroimaging studies showed reduced self-touch-evoked activity in the somatosensory cortex and cerebellum compared to identical external or delayed self-touch (Blakemore et al., 1998; Hesse et al., 2010; Kilteni et al., 2023; Kilteni & Ehrsson, 2020).

If attenuation is driven by forward-model prediction, neural correlates should emerge *before* sensory input, yet evidence for pre-stimulus predictive activity remains elusive. Previous studies have focused on stimulus-evoked activity (Hesse et al., 2010), or averaged across periods containing movement and touch (Blakemore et al., 1998; Kilteni & Ehrsson, 2020), making anticipatory and post-stimulus activity difficult to disentangle. Two fMRI studies reported increased connectivity between somatosensory cortices and the cerebellum during self-touch relative to external or delayed self-touch (Kilteni et al., 2023; Kilteni & Ehrsson, 2020), but could not determine neither whether this connectivity increased before the touch, nor the directionality (i.e., cerebellar predictions to cortex or somatosensory input projected to the cerebellum).

To resolve these issues, we recorded magnetoencephalography (MEG) while participants produced a self-touch on their left index finger with their right index finger. Neural activity was compared with an external touch condition (identical touch without movement) and a misaligned condition, in which participants made the same movement and received the same touch, but with the hands spatially misaligned to disrupt sensory predictions. Thus, the misaligned condition included a self-initiated touch, but reduced the correspondence between the efference copy, limb configuration, and stimulated body site. In other words, although participants could learn that moving one hand causes touch on the other hand, the misaligned configuration was expected to support a less precise or less canonical prediction of touch at that body site than the spatially aligned self-touch condition. Conditions were kept short to limit extensive learning or recalibration of the misaligned mapping. Our misaligned and self-touch conditions, therefore, targeted differences in the strength of spatially specific predictions, rather than a simple contrast between “predicted” and “unpredicted” touch. The misaligned condition further controlled for differences between self-touch and external touch, including simultaneous movement and touch, bilateral tactile stimulation, temporal predictability, divided attention, and anticipation (Bays & Wolpert, 2008; Kilteni & Ehrsson, 2017, 2020).

We included no-touch blocks for all conditions to control for motor, postural, proprioceptive, and visual differences. Subtracting stimulus-absent from stimulus-present trials within each condition isolates activity related to the tactile consequence of the action. Finally, participants completed a force discrimination task after MEG to quantify attenuation under behaviourally matched conditions. (Kilteni, 2023)

We hypothesised that self-touch would elicit reduced somatosensory-evoked activity compared to external and misaligned touch. We further expected pre-stimulus beta-band (13-30 Hz) modulations over contralateral sensorimotor cortex to index predictive processes (Andersen & Dalal, 2021, 2024; Andersen & Lundqvist, 2019; Kimura, 2021; Pando-Naude & Andersen, 2025; van Ede et al., 2010, 2011a), particularly differences between self-touch and misaligned touch, which share temporal expectations but differ in spatial specificity. Finally, we predicted increased pre-stimulus beta-band connectivity from left cerebellar lobule VI to the somatosensory cortex during self-touch, motivated by fMRI evidence of cerebellar involvement in self-touch attenuation (Kilteni & Ehrsson, 2020), along with stronger perceptual attenuation for self-touch than for external or misaligned touch.

## Materials and Methods

### Participants

Twenty-four adults completed the experiment (aged 19-36 years; 12 female, 21 right-handed, 3 ambidextrous). Two participants were replaced, one who did not attend the MRI session and one who did not attend the MEG session (a total of 26 participants recruited). Exclusion criteria were reports of any current or history of psychological or neurological conditions, as well as the use of any psychoactive drugs or medication to treat such conditions. Handedness was assessed using the Edinburgh Handedness Inventory (Oldfield, 1971) with an average handedness score of 75.48 (*SD* = 34.56). All experiments were approved by the Swedish Ethical Review Authority (registration no. 2021-02319). All participants provided written informed consent and were compensated for their time.

### Structural MRI data acquisition

Structural MRI data were acquired from each participant to co-register with the MEG data for source reconstruction. 3D T1-weighted magnetisation prepared rapid gradient echo (MPRAGE) sequence structural images (voxel size: 1x1x1 mm) were obtained on a Siemens Prisma 3.0 T MR scanner. The pulse sequence parameters were: 1 mm isotropic resolution; field of view: 256 x 256 mm; 208 slices; slice thickness: 1 mm; bandwidth per pixel: 240 Hz/pixel; flip angle: 9°; inversion time (TI): 900 ms; echo time (TE): 2.98 ms; repetition time (TR): 2300 ms.

### Preparation of participants

Before MEG measurement, each participant’s head shape was digitized using a Polhemus FASTRAK. Three fiducial points (the nasion, the left and right pre-auricular points) were digitized along with the positions of four head-position indicator coils (HPI-coils). Approximately 200 extra points were digitized over the head shape of each participant.

### Acquisition of MEG data

MEG data were recorded with an Elekta Neuromag TRIUX 306-channel MEG system, with 102 magnetometers and 204 planar gradiometers. The MEG scanner was located inside a two-layer magnetically shielded room. Data were sampled at 1000 Hz with an online 0.1 Hz high-pass filter and 330 Hz low-pass filter.

### MEG stimuli, apparatus G task

Tactile stimuli were generated using an inflatable membrane (MEG International Services Ltd., Coquitlam, Canada) attached to the left index fingertip. The membrane was part of a custom stimulation rig controlled by pneumatic valves (model SYJ712M-SMU-01F-Q, SMC Corporation, Tokyo, Japan) using 1 bar of pressurised air.

**Figure 1a** illustrates the experimental setup. **Figure 1b** shows the trial procedure for each condition. In the external *touch* condition, each trial began with the presentation of a blink instruction (the word “blink” in the centre of the screen). After 1200 ms, the blink instruction was replaced with a fixation cross cue that appeared in the centre of the screen. After 500 ms, the tactile stimulus of 100 ms duration was delivered to the pulp of the left index finger. Following the stimulation, an inter-trial interval was presented, which randomly varied in duration between 800, 1000, and 1200 ms. The fixation cross remained on the screen for the duration of the inter-trial interval before it was replaced with the blink instruction of the following trial. The inter-stimulus interval was therefore 2800 ± 200 ms. Participants were instructed to rest both hands during the external touch block. The left hand rested palm-up and its position was fixed using a vacuum pillow (Germa Vaccum Pillow).

**Figure 1.**
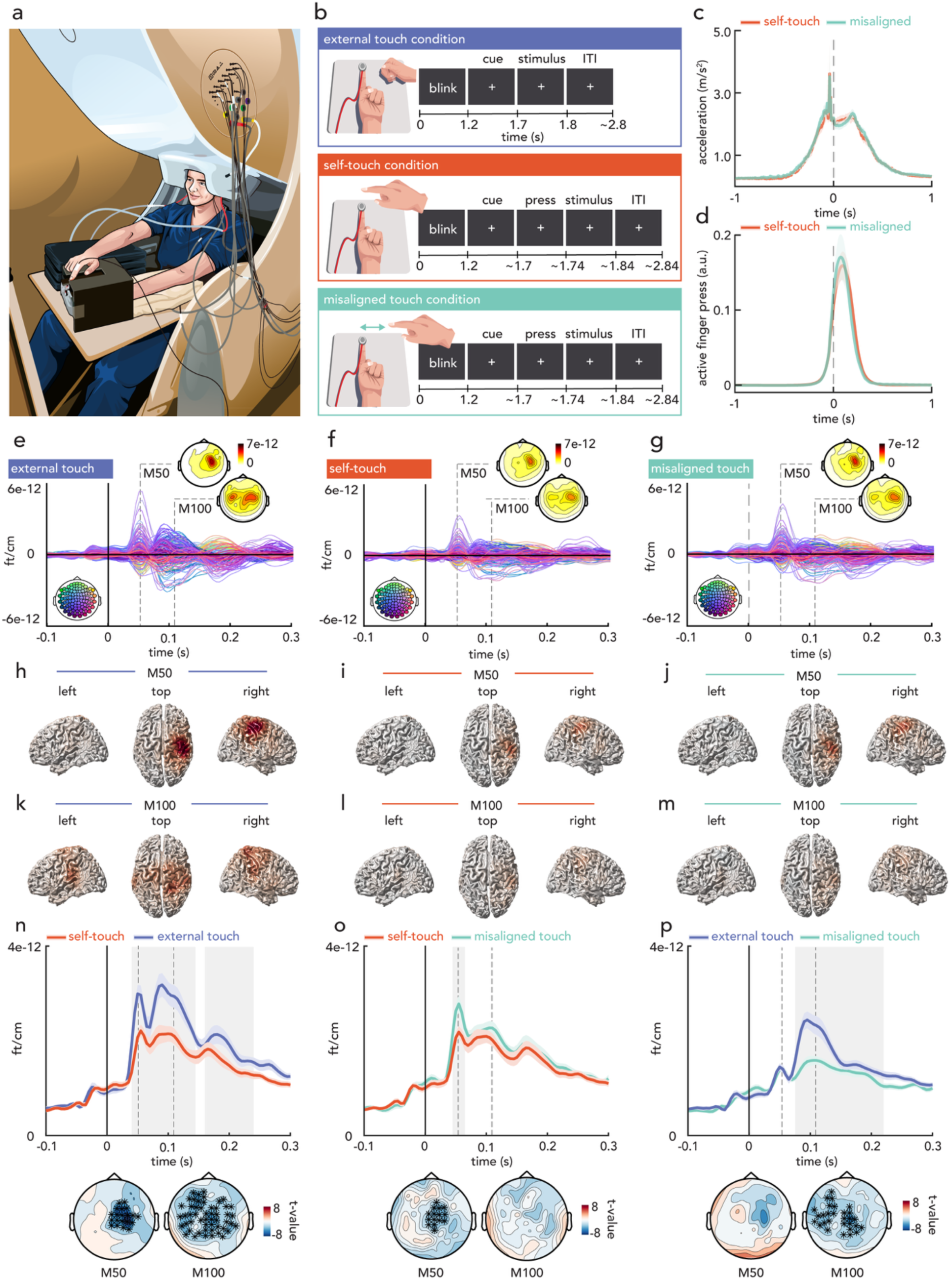
MEG stimulus-evoked results. **(a)** Illustration of the MEG setup, showing the custom device used for simulating self-generated touch. Participants pressed a lever with their right index finger, triggering a tactile stimulus on their left index finger. **(b)** Experimental conditions. External touch: The touch was delivered automatically after the visual cue. Self-touch: The touch was delivered when participants pressed a lever above their left index finger. Misaligned touch: Identical to the self-touch condition, except that the device was positioned 25 cm to the right of the left index finger. **(c)** The conditions with movement (self-touch and misaligned touch) did not significantly differ in their acceleration profiles for the moving hand or **(d)** the force exerted on the response device that triggered the tactile stimulus (see **Supplementary Text S1** for details). Shaded error bands show ± standard error of the mean (s.e.m.). **(e, f, g)** MEG sensor level results. Butterfly plots of the 204 gradiometers, time-locked to the onset of the tactile stimulus on the left index finger, with topographical plots of the root mean square values of combined gradiometer pairs. Topographies show the M50 (50 ms) and M100 (110 ms) components of the event-related fields. Data are presented as the difference between stimulus-present and stimulus-absent blocks (see **Supplementary Figure S1c**). Coloured heads in the bottom-left corners indicate sensor locations. **(h, i, j)** Linearly constrained minimum variance (LCMV) beamformer source reconstructions of the M50 component (45-65 ms) and **(k, l, m)** the M100 component (90-120 ms). **(n, o, p)** Grand averaged responses of the combined gradiometer pairs for each statistical comparison, with shaded error bands representing ± s.e.m. The grey areas mark the clusters that informed the rejection of the null-hypothesis in the cluster-based permutation tests. The topographical plots show the uncorrected mass-univariate t-values at the M50 and M100 time points, with cluster channels highlighted. Overall, the figure shows that the M50 response was significantly attenuated for the self-touch compared to external touch and misaligned touch conditions. Significant differences in the external touch vs. misaligned touch comparison emerged later (> 75 ms), peaking around the M100 component.

In the self-touch condition, the same blink instruction and fixation cross cue were presented on each trial. Participants were instructed to tap with their right index finger on a custom-made lever device placed directly above, but not in contact with, their left index finger when the fixation cross cue appeared. Participants had a maximum window of 3000 ms to press the lever. When the lever device was pressed with the right index finger, an optical fibre placed below the device detected the proximity of the finger, which triggered the onset of the tactile stimulus on the left index finger. The setup simulated pressing the left index finger with the right index finger through an object. Participants were instructed to keep their right index finger on a start position marker 5 cm from the lever device before and after the pressing action. The misaligned touch condition was identical to the self-touch condition, except that the device and start-position marker were placed 25 cm to the right of the left hand. Thus, the action of moving 5 cm to press the device in the self-touch and misaligned touch conditions was identical, but the spatial configuration no longer simulated touching one’s left index finger. The inter-stimulus interval did not significantly differ between conditions (see **Supplementary Text S1**). The average movement onset was 418 ms before stimulus onset (SD = 215) for the self-touch condition and 404 ms (SD = 160) for the misaligned touch condition, and no significant difference was found (see **Supplementary Text S1**).

Participants completed 2 blocks of each condition. Each block consisted of 90 trials, resulting in a total of 180 trials per condition (self-touch, external touch and misaligned touch). Each condition was also completed in a ‘stimulus-absent’ block in which no stimulus was delivered. In these stimulus-absent blocks, the same blink instruction and fixation cross were presented on each trial, but no stimulus was delivered. Participants were given self-timed breaks between blocks and instructions were presented on the screen before each block. The task lasted approximately 1 hour. The order of the conditions (external touch, self-touch, and misaligned touch) was counterbalanced across participants. The order of the stimulus-present and stimulus-absent blocks was randomised across participants, such that 50% of participants started with stimulus-present blocks and 50% of participants started with stimulus-absent blocks. Thus, for half of the participants, the first block of a given condition was stimulus-present, and for the other half, it was stimulus-absent. The blocks then alternated throughout the session so that each condition (self/misaligned/external) and stimulation block (present/absent) was distributed across the whole recording period.

White noise was played through sound tubes (model ADU1c, KAR Oy, Helsinki, Finland) into the ears of participants at approximately 65 dB throughout the task, making the tactile stimulation inaudible. An accelerometer, attached to the dorsal surface of the middle phalanx of the right index finger, measured the acceleration of the finger movements along three orthogonal axes. The continuous time course of the accelerometer was sampled with the MEG data and averaged offline by calculating the Euclidean norm.

### MEG Data processing

#### Pre-processing

MEG data were pre-processed offline by first applying temporal signal space separation (tSSS) to suppress artefacts from outside the scanner helmet and correct for head movement during the recordings (Taulu et al., 2004; Taulu & Simola, 2006). Head origin was shifted to a position based on the initial position for each participant.

The data analysis was performed using the FieldTrip toolbox for EEG/MEG-analysis (Oostenveld et al., 2011); Donders Institute for Brain, Cognition and Behaviour, Radboud University, the Netherlands. See http://fieldtriptoolbox.org). The data was segmented into 3-second epochs from 1.5 seconds before stimulus onset to 1.5 seconds after stimulus onset for the stimulus-present blocks. For stimulus-absent blocks, the data were also segmented into 3-second epochs centred around a ‘dummy’ stimulus delivered at the same timepoint within the trial as the stimulus was presented in stimulus-present blocks. The data were down-sampled to 200 Hz and demeaned using the entire epoch. The delay between the digital trigger and the onset of the stimulation was assessed to be 41.0 ms via a separate recording using an accelerometer attached to the inflatable membrane. This delay was subtracted from each epoch. Data epochs were cleaned semi-manually using the *ft_rejectvisual* function, with segments showing large variances removed without applying a fixed threshold. The method was applied blind to the experimental condition. On average, 46 (SD = 83.01) trials were rejected per participant (< 5% of total trials). An ANOVA with a factor of condition (self-touch, misaligned touch and external touch) and stimulation (stimulus-present and stimulus-absent) was run on the number of rejected trials, for which no significant main effects or interactions were observed (all p-values > 0.05). To attenuate artefacts (e.g. cardiac activity, eye blinks and movements), we applied Independent Component Analysis (ICA). The data were decomposed into 60 independent components per participant using the *runica* algorithm (Infomax ICA). The resulting components were inspected visually using sensor-space topographies and time courses. An average of 9.0 components (SD = 6.3) were projected out of the data using linear back-projection.

#### Stimulus-evoked activity

Event-related Fields (ERFs) were extracted by averaging the trials from each of the three stimulus-present blocks (external, self, and misaligned) as well as the trials from the corresponding stimulus-absent blocks (external, self, and misaligned). The averaged stimulus-evoked waveforms were then subtracted from the averaged waveforms of the corresponding stimulus-absent conditions. This subtraction minimises non-stimulus-related activity, making the conditions more comparable. Importantly, this minimises any influence of posture, generic arousal, motivation or motor-preparation differences. The subtraction resulted in three conditions of interest (self-touch, external touch, and misaligned touch).

#### Source reconstruction of the M50 ERF component

The time course of neural activity within regions of interest was reconstructed using a Linearly Constrained Minimum Variance (LCMV) beamformer technique (Van Veen et al., 1997). T1-weighted MRI images were segmented and resliced to align with the headpoints obtained for each participant during the MEG preparation. A single-shell head model was computed based on the segmented MRI (Nolte, 2003). A standard source model with a 10 mm grid, aligned to MNI space, was used for source reconstruction. The source model was registered to each participant’s MRI. A common spatial filter was calculated based on the data covariance matrix computed over a 0–200 ms post-stimulus window, using combined trials from each stimulus-present condition and its respective stimulus-absent condition. A regularisation parameter (lambda) of 5% was applied to the covariance matrix. The common spatial filter was then applied to the averaged evoked waveforms for each condition. A fixed orientation was determined using single value decomposition (SVD), and the dipole moment was projected onto this axis. Source strength was quantified by averaging the projected dipole moment over a latency window, either 45-65 ms (M50 component) or 90-120 ms (M100 component). Finally, power in the stimulus-present condition was then expressed relative to the stimulus-absent condition (e.g., external = (external stimulus present – external stimulus absent) / external stimulus absent).

#### Pre-stimulus time-frequency activity

For induced activity, the time-locked response of each condition was subtracted from each segment of the corresponding condition. Subtracting the part of the signal that reflects phase-locked or evoked components prior to computing time-frequency representations (TFRs) minimises the presence of time-locked activity in the induced activity (Kalcher & Pfurtscheller, 1995; Pfurtscheller & Lopes Da Silva, 1999). TFRs of individual trials were then calculated using Morlet wavelet analysis with a wavelet width that linearly increased from 3 to 18 across a frequency range of 1-80 Hz. The planar gradient magnitude over both directions was computed by summing the two gradients at each sensor to a single positive-valued number. The averaged TFRs for the stimulus-present conditions were subtracted from the averaged TFRs of the corresponding stimulus-absent conditions.

#### Source reconstruction of pre-stimulus beta

Source reconstruction was performed using the Dynamic Imaging of Coherent Sources (DICS) (Gross et al., 2001) beamforming method in FieldTrip (Oostenveld et al., 2011). Cross-spectral density (CSD) matrices were computed for each condition over the 13-30 Hz frequency band in the pre-stimulus window (-500ms 0 ms). A common spatial filter was computed from the combined data of each stimulus-present and stimulus-absent condition. A regularisation parameter (lambda) of 5% was applied to the CSD matrix. Source power was then estimated separately for each condition using this common filter. Finally, contrasts were calculated by expressing power in the stimulus-present condition relative to the stimulus-absent condition.

#### Calculation of virtual sensors

Virtual channel time series were estimated based on linearly constrained minimum variance beamformer (LCMV) projections to two ROIs derived from previous fMRI studies investigating self-touch attenuation when the right hand moves to touch the left hand. The right primary somatosensory cortex (MNI: x = 48, y = -18, z = 60) was chosen based on Kilteni et al. (2023) and the left cerebellar lobule VI (MNI: x = -22, y = -58, z = -22) was chosen based on Kilteni and Ehrsson (2020). The template coordinates were transformed to individual coordinates.

#### Coherence analysis

Spectral coherence between the virtual sensors was computed using FieldTrip’s *ft_connectivityanalysis,* retaining only the imaginary component of coherence (‘*absimag*’ option) to minimise spurious zero-lag correlations. Time-frequency representations (TFRs) of power and cross-spectral density were computed using multitaper convolution with a Hanning taper. Frequency-dependent time windows of 7 cycles per frequency were used. Coherence estimates were calculated for each condition in the beta band (13-30 Hz), separately for each time point in the pre-stimulus window (-1 to 0 seconds). For each condition, the stimulus-absent block was subtracted from the corresponding stimulus-present block to isolate stimulus-specific coherence.

#### Granger causality analysis

Directed connectivity between the virtual sensors was assessed using non-parametric Granger causality using FieldTrip’s *ft_connectivityanalysis* function with the ‘*granger*’ method and ‘*bivariate*’ spectral factorisation. To improve signal stationarity, the analysis was confined to a 500 ms pre-stimulus interval (-0.5 to 0 ms) that was closest to stimulus onset. Frequency decomposition was performed using the multitaper method with ±2 Hz spectral smoothing. For each participant and condition, stimulus-present Granger causality values were expressed relative to stimulus-absent values. Time-reversed trials were also analysed as a control to ensure directional asymmetries were not due to spurious temporal correlations or non-causal dependencies. Additional analyses of direction-reversed connectivity (from S1 to the cerebellum) were conducted to assess the directional specificity.

#### Behavioural force-discrimination task

Following MEG data collection, participants completed the force discrimination task (Kilteni, 2023). They rested their left hand, palm up, with their index finger placed on a moulded support (Figure 3a). On each trial, a motor (Maxon EC Motor EC 90 flat; Switzerland) delivered two forces (the *test* force and the *comparison* force) on the pulp of the left index finger through a cylindrical probe (25 mm height) with a flat aluminium surface (20 mm diameter) attached to a lever on the motor. A force sensor (FSG15N1A, Honeywell Inc.; diameter, 5 mm; minimum resolution, 0.01 N; response time, 1 ms; measurement range, 0–15 N) within the probe recorded the forces applied on the left index finger. Following the presentation of the two forces, participants verbally reported which of the two forces felt stronger, the first or the second. A second identical force sensor within an identical cylindrical probe (“active force sensor”) was placed on top of, but not in contact with, the probe of the left index finger.

Participants judged the intensity of a *test* force and a *comparison* force (100 ms duration each), separated by a random interval between 800 ms and 1200 ms in a two-interval forced-choice (2IFC) task. The intensity of the *test* force was 2 N, while the intensity of the *comparison* force was systematically varied among seven force levels (1, 1.5, 1.75, 2, 2.25, 2.5, or 3 N). In all the conditions, the forces were delivered by the same motor, in order to precisely control their magnitude, however, the source of the force was manipulated across conditions such that the force was triggered by the participant’s contact with a force sensor (self-touch condition and misaligned touch condition) or automatically by the stimulus computer (external touch condition).

In the external touch condition, participants rested both hands and received the test and the comparison force on the left index finger. This external condition was used to assess somatosensory perception in the absence of any movement (Bays et al., 2005; Kilteni et al., 2019, 2020). Each trial began with an auditory cue (100 ms duration, 997 Hz) followed by the *test* force delivered to the participant’s left index finger 800 ms after the cue by the motor. The *comparison* force was then delivered at a random interval between 800 ms and 1200 ms after the *test* force.

In the self-touch condition, the same auditory cue was presented as in the external touch condition, but participants now actively tapped on the force sensor placed on top of, but not in contact with, the probe. The active tap on the force sensor triggered the motor to apply the *test* force on the left index finger (threshold 0.2 N). The tap of their right index finger triggered the *test* force on their left index finger with an intrinsic delay of 36 ms. The *comparison* force was then delivered at a random interval between 800ms and 1200ms after the *test* force. Participants were asked to tap, neither too weakly nor too strongly, with their right index finger, “as if tapping the screen of their smartphone”, as instructed in previous studies (Asimakidou et al., 2022; Kilteni et al., 2021; Kilteni C Ehrsson, 2022).

The misaligned touch condition was identical to the self-touch condition, except that the force sensor that participants tapped to trigger the touch on their left index finger was placed 25 cm to the right of the left index finger, rather than directly above the left index finger. While previous studies investigating spatial manipulations of self-touch have used force-matching paradigms (Bays C Wolpert, 2008; Kilteni C Ehrsson, 2017, 2020). These typically involve sustained force stimuli lasting several seconds, unlike the brief tactile stimuli used in the MEG task. By contrast, the force discrimination task used stimuli of identical duration to those in the MEG task, enabling a more closely matched behavioural task. To our knowledge, this is the first study to apply a spatial manipulation of self-touch within the force discrimination task.

White noise was played throughout the task to mask any sounds made by the motor that could serve as a cue for the task. Each block consisted of 70 trials, resulting in 210 trials per participant. The order of the conditions (self-touch, external touch, and misaligned touch) was counterbalanced across participants.

#### Preprocessing of psychophysical trials

The behavioural task included 5040 trials in total (24 participants * 70 trials * 3 conditions). 20 trials were excluded because of 16 missing responses and 4 trials in which the force was not applied correctly (1.85 N < *test* force < 2.15 N)

#### Fitting of psychophysical responses

For each experiment and each condition, the responses were fitted with a generalized linear model using a *logit* link function (Equation 1):

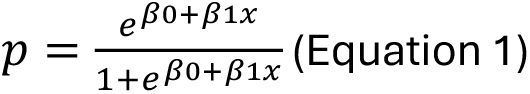

We extracted two parameters of interest: the Point of Subjective Equality (PSE) 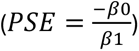, which represents the intensity at which the *test* force is felt as strong as the *comparison* force (*p* = 0.5) and quantifies the perceived intensity, and the JND 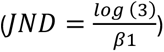 which reflects the participants’ discrimination ability. Before fitting the responses, the values of the applied *comparison* forces were binned to the closest value to their theoretical values (1, 1.5, 1.75, 2, 2.25, 2.5, or 3 N). See **Supplementary Figure S3a** for individual participant psychometric curves. The fitted logistic models were very good, with McFadden’s R- squared measures ranging between 0.600 to 0.970.

After the psychophysical task, participants completed the Schizotypal Personality Questionnaire (SPQ) (Raine, 1991) which addressed a separate research question and will be reported in a different manuscript.

### Statistical analyses

For movement-related data from the MEG session (i.e., movement onsets, maximum acceleration, and press intensity), repeated measures analysis of variance (rmANOVA) was used with two factors: condition (self-touch vs. misaligned touch) and stimulation block (stimulus-present vs. stimulus-absent). For behavioural data generated from the force discrimination task (i.e. PSEs and JNDs), rmANOVA was used to compare the external touch, self-touch, and misaligned touch conditions. For the peak force exerted with the right index finger on the force sensor, a paired t-test was used to compare the self-touch and misaligned touch conditions. Significant main effects from the rmANOVAs were followed by Bonferroni-corrected post hoc comparisons. Effect sizes are reported as partial eta squared 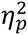 for F-tests and Cohen’s *d* for t-tests. 95% confidence intervals (*CI^S5^*) are reported for each statistical test. All tests were two-tailed.

For stimulus-evoked activity, comparisons were made between the self-touch and external touch conditions, as well as self-touch and misaligned touch, and misaligned touch and external touch conditions. For pre-stimulus activity, only conditions that involved action were compared (i.e., self-touch and misaligned touch), given that comparisons between a condition that involves action and the external touch condition without action would likely reflect the presence of movement within the pre-stimulus window, or activity locked to the fixation cross cue. Non-parametric cluster permutation (Maris & Oostenveld, 2007) was used to compare the conditions. The following steps are taken for significance testing with cluster-based permutation tests: 1) perform mass-univariate dependent samples t-statistics comparing each condition for each of the samples in the multidimensional data structure; 2) neighbouring data points in the multidimensional data structure below a threshold of *p* < 0.05 are summed to calculate the cluster level statistic; 3) the procedure is repeated for 2000 permutations with the condition labels shuffled on each permutation; 4) The maximum cluster statistic was evaluated under its permutation distribution (shuffled data). The cluster-level significance threshold was set at the two-tailed level of 0.025. For the analysis of stimulus-evoked activity, the test window spanned 0 to 400 ms post-stimulus. For the analysis of pre-stimulus oscillatory activity, the test window covered -1000 to 0 ms relative to stimulus onset and was restricted to the beta band (13-30 Hz).

## Results

### Decreased somatosensory cortical activity evoked by self-touch

We first investigated attenuation of somatosensory activity time-locked to the touch. In the MEG session (Figure 1a), participants received tactile stimuli in three conditions (external touch, self-touch, and misaligned touch) (Figure 1b). Control analyses showed no significant effects of condition on the movement onset, movement acceleration (Figure 1c), press intensity (i.e., how much participants pressed on the lever with their moving hand) (Figure 1d), or the inter-press-interval when comparing the self-touch and misaligned touch conditions (**Supplementary Text S1**). Therefore, any differences in brain activity between the movement conditions are unlikely to be driven by systematic differences in movement kinematics.

Event-related fields (ERFs) (**Figure 1e-g**) were computed by averaging trials in each of the three conditions and subtracting the stimulus-absent waveforms, minimising movement-related activity and any postural or visual differences (See *Methods – Stimulus-evoked activity &* **Supplementary Figure S1c**). The somatosensory M50 and the M100 components of the ERF were observed, as expected. The M50 component was source localised to the right postcentral gyrus (i.e., primary somatosensory cortex contralateral to the stimulated finger) (**Figure 1h-j**). The M100 component was localised to a broader area including bilateral sensorimotor areas and bilateral parietal operculum (i.e., bilateral secondary somatosensory cortices) (**Figure 1k-m**). Thus, the source localisation confirms previous findings that the M50 and M100 components reflect activity in primary and secondary somatosensory cortices, respectively (Hesse et al., 2010; Yamashiro et al., 2019).

Cluster-based permutation tests (Maris & Oostenveld, 2007) showed a significant difference between the self-touch condition and the external touch condition (*p* < 0.001) (**Figure 1n**). The first cluster that informed the rejection of the null hypothesis indicated lower amplitudes in the self-touch condition compared to the external touch condition from 40 ms to 145 ms, including the M50 and M100 components and a later second cluster between 160 ms and 240 ms. Cluster-based permutation tests also showed a significant difference (*p* = 0.015) when comparing the self-touch with the misaligned touch condition (**Figure 1o**), indicating lower amplitudes in the self-touch compared to the misaligned touch condition between 45 ms and 65 ms, corresponding to the M50 component. Finally, cluster-based permutation tests showed a significant difference between the misaligned touch and external touch condition (*p* < 0.001) (**Figure 1p**), indicating lower amplitudes in the misaligned touch condition compared to the external touch condition between 75 ms and 160 ms corresponding to the M100 component, and between 155 ms and 220 ms.

Together, these results show that both early (M50) and late (M100) somatosensory activity were attenuated for self-touch compared to externally generated touch, even though the stimuli were physically identical. Importantly, when comparing the self-touch and the misaligned touch conditions, which were matched in self-initiation and action-touch contingency but differed in spatial sensorimotor congruence, attenuation of somatosensory activity was specific to the earlier M50 component. Later components were similarly attenuated in the self-touch and misaligned touch conditions relative to the external touch condition, likely reflecting additional effects between movement and non-movement conditions, including temporal predictability of touch, feelings of agency, and general anticipation of touch.

### Greater pre-stimulus beta desynchronization preceding self-touch

To avoid confounding movement-related activity with predictive effects, we focused our pre-stimulus analysis on the self-touch and misaligned touch conditions, excluding the external touch condition. Notably, comparisons with the external touch condition (which lacked movement and had a fixed cue timing) gave similar results, but they are reported separately in the **Supplementary Text S2** due to inherent differences in motor involvement and cue-locked activity.

To test whether predictive mechanisms engage before self-touch, we analysed pre-stimulus (-1000 to 0 ms) beta-band (13-30 Hz) oscillatory activity in the self-touch and misaligned touch conditions (**Figure 2a-c**). Identically to the touch-evoked analyses, we first subtracted the activity from the stimulus-absent blocks (see **Supplementary Figure S2a**). Source localisation identified the origin of the pre-stimulus beta desynchronisation in the sensorimotor cortex contralateral to the stimulated finger (**Figure 2d**).

**Figure 2.**
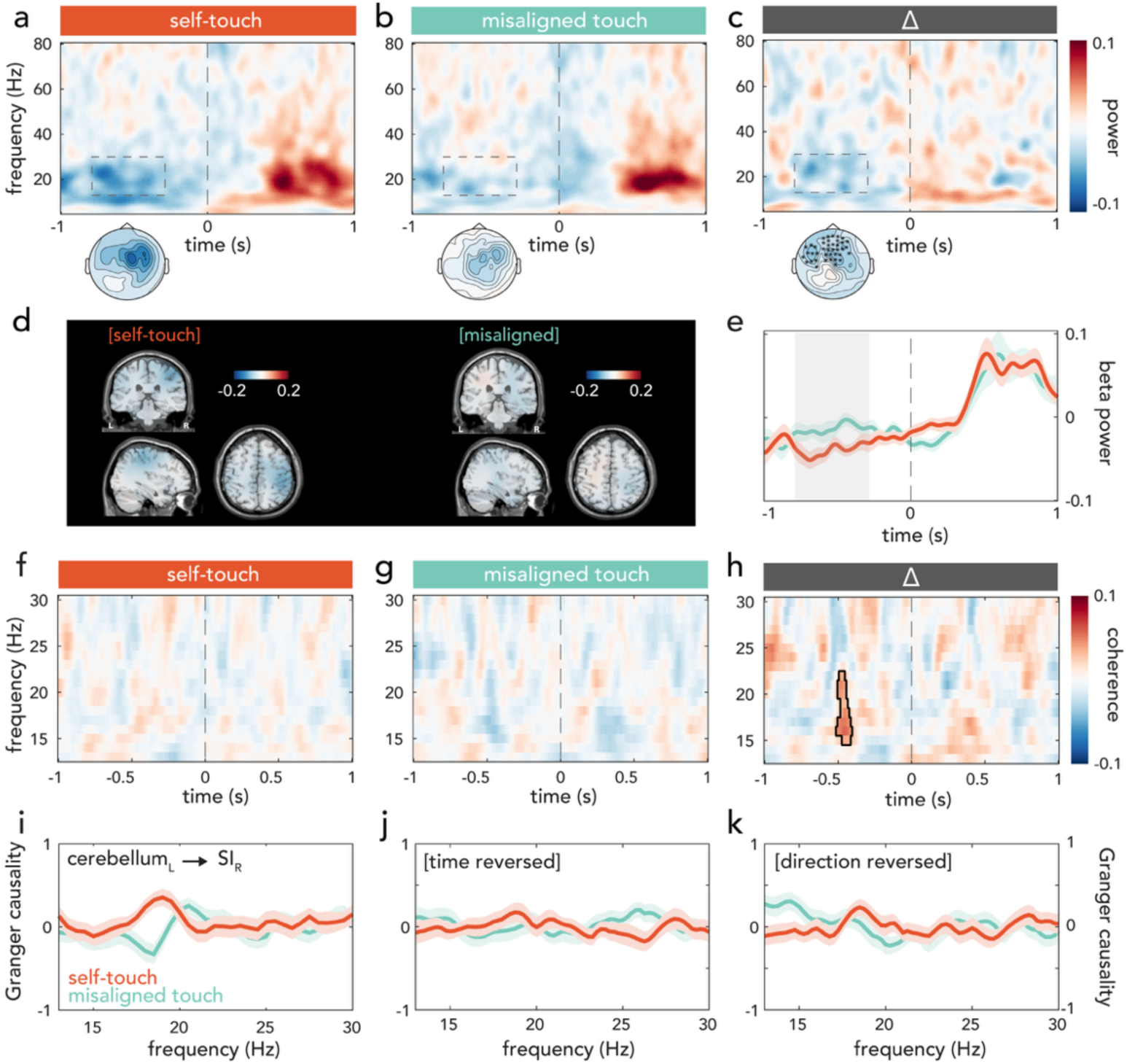
MEG pre-stimulus results. **(a-c)** Grand-averaged time-frequency power over channels contributing to the cluster showing a significant difference between conditions (self-touch vs. misaligned touch): **(a)** the self-touch condition, **(b)** the misaligned touch condition and **(c)** their difference (Δ = self-touch - misaligned touch). Stronger beta-band (13-30 Hz) desynchronisation was found in the self-touch condition before stimulus onset (dashed box). Topographies show the average pre-stimulus beta power with cluster channels highlighted. **(d)** Source localization of pre-stimulus beta activity revealed sensorimotor sources. **(e)** Line plot of mean beta power across time, averaged over the channels contributing to the cluster informing the rejection of the null hypothesis of no difference between conditions. Time points contributing to the cluster-level effect are shown in grey. **(f-h)** Grand averaged beta-band coherence differences (stimulus present minus stimulus absent) between the right primary somatosensory cortex (SIR) and left cerebellum for **(f)** the self-touch condition, **(g)** the misaligned touch condition, and **(h)** their difference (Δ = self-touch – misaligned touch). The outlined region marks the cluster that informs the rejection of the null hypothesis. **(I, j, k)** Non-parametric Granger causality estimates of directed connectivity in the beta-band. **(i)** Connectivity in the pre-stimulus window (-0.5 to 0 ms) from the left cerebellar lobule VI to the right primary somatosensory cortex was significantly higher before self-touch compared to misaligned touch. Control analyses found no significant differences when **(j)** reversing the time axis and **(k)** reversing the direction (SIR to cerebellumL). Shaded error bands show ± standard error of the mean (s.e.m).

**Figure 3.**
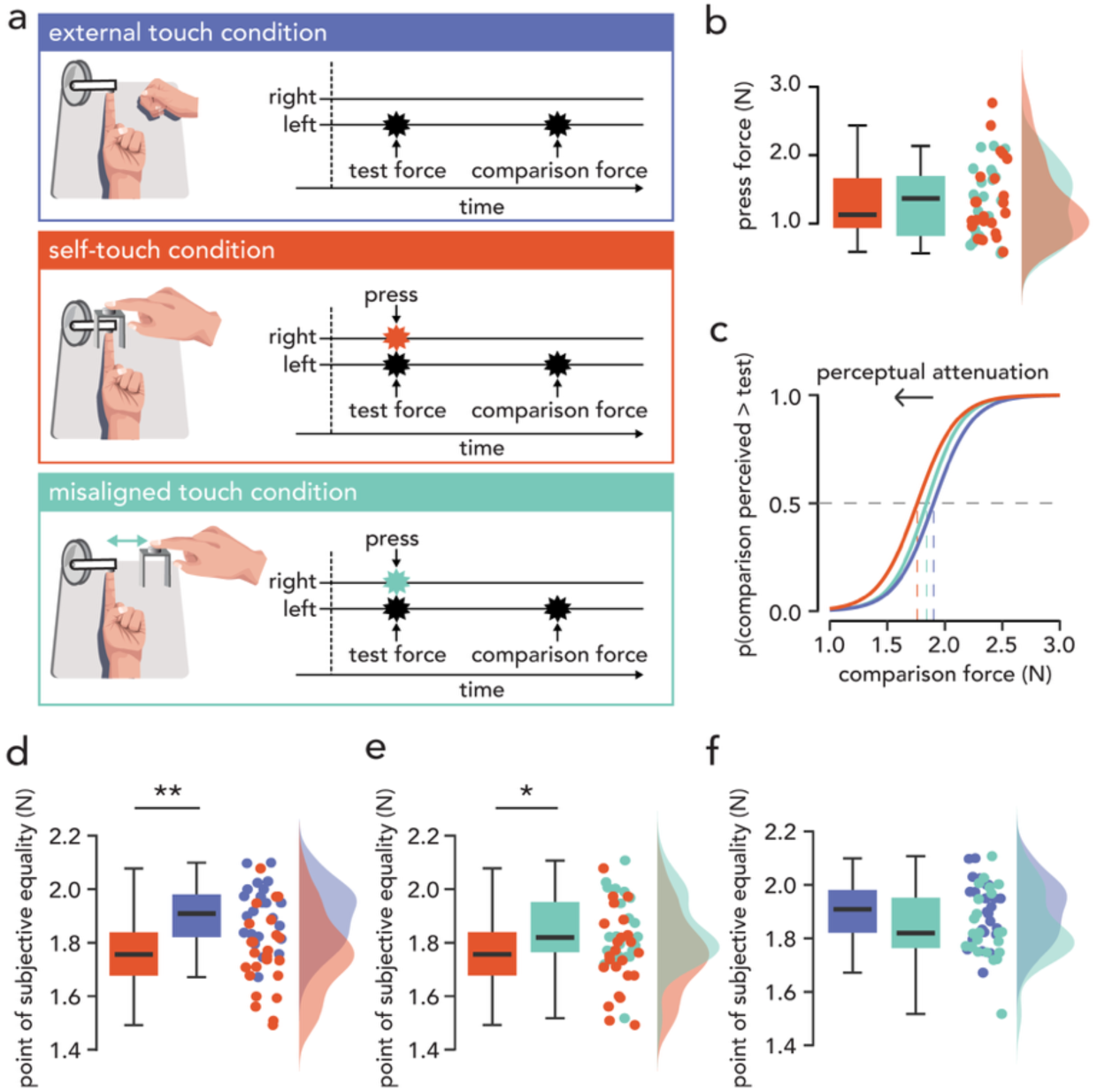
Behavioural results. **(a)** Experimental conditions and trial procedure for the force discrimination task. In all three conditions, participants discriminated the intensity of a test force (2 N) from a comparison force (1-3 N) delivered to their left index finger on each trial. External touch: The test force was delivered automatically after the onset of an auditory cue on each trial. Self-touch: The test force was delivered when participants pressed a force sensor above their left index finger with their right index finger. Misaligned touch: Identical to the self-touch condition, except that the force sensor was positioned 25 cm to the right of the left index finger. **(b)** Participants pressed with approximately equal force in the self-touch and misaligned touch conditions. **(c)** Group psychometric curves for each condition using the average JND and PSE values across participants. The leftward shift in the self-touch condition indicates attenuated PSE values compared to the misaligned touch and external touch conditions. **(d)** Self-touch was perceived as weaker than external touch and **(e)** misaligned touch, whereas **(f)** external and misaligned touch did not significantly differ in perceived intensity. **(b, d-f)** Boxplots show the median and interquartile range. Raincloud plots show the individual participant values (dots) and their distributions. **p<0.001, *p<0.05.

A cluster-based permutation test on the sensor-level data showed a significant difference between the self-touch condition and the misaligned touch condition (*p* = 0.0245), confirming reduced beta power in the self-touch condition compared to the misaligned touch condition in the pre-stimulus window (**Figure 2e**). This finding suggests that when self-generated touch occurs at a congruent body location, the brain’s prediction of the sensory consequence of the action is associated with a desynchronisation of beta oscillations over sensorimotor cortex before the touch occurs. We interpret these pre-stimulus beta modulations as reflecting differences in the relative strength of spatially specific predictions about self-generated touch.

### Increased cerebellar-somatosensory connectivity preceding self-touch

Having established condition-related differences in pre-stimulus beta desynchrony, we next examined changes in functional connectivity between two regions of interest (ROIs). For the cerebellum we focused on the left cerebellar lobule VI, based on earlier anatomical evidence that the cerebellum controls output to, and receives input from, the ipsilateral side of the body (Grodd et al., 2001; Manni & Petrosini, 2004; Mottolese et al., 2013) and earlier fMRI evidence that this region is involved in self-touch attenuation when the right hand moves to touch the left hand (Kilteni & Ehrsson, 2020). Although deeper structures like the cerebellum pose challenges for MEG recordings, recent simulations and empirical studies have demonstrated that cerebellar activity can be reliably detected with MEG (Andersen et al., 2020; Samuelsson et al., 2020). We tested all these hypotheses with MEG because its millisecond temporal resolution allows dissociation of pre- and post-stimulus neural activity, crucial for identifying predictive processes. Two ROIs were derived from previous fMRI studies investigating self-touch attenuation when the right hand moves to touch the left hand. The right primary somatosensory cortex (MNI: x = 48, y = -18, z = 60) was chosen based on Kilteni et al. (2023) and the left cerebellar lobule VI (MNI: x = - 22, y = -58, z = -22) was chosen based on Kilteni and Ehrsson (2020). Identically to all previous analyses, we first subtracted the activity from the stimulus-absent blocks to account for any movement, posture or visual differences between the conditions.

A cluster-based permutation test on the coherence time course indicated significantly higher coherence in the beta band (13-30 Hz) between right SI and left cerebellum for the self-touch condition compared to the misaligned touch condition (*p* = 0.019) (**Figure 2f-h**). Coherence in the stimulus-present and stimulus-absent blocks is shown separately in **Supplementary Figure S2b** and statistical comparisons are provided in Supplementary Text S3.

To further characterise the nature of the cerebellar-somatosensory connectivity, we conducted a hypothesis-driven Granger causality analysis restricted to the right SI and the left cerebellar lobule VI in the pre-stimulus window. ROIs were selected based on the areas identified with fMRI (Kilteni et al., 2023; Kilteni & Ehrsson, 2020). Granger causality provides complementary information to coherence by estimating the direction of influence between regions. A cluster-based permutation test on the Granger causality values confirmed significantly higher directed connectivity in the beta band (13-30 Hz) from the left cerebellum to the right SI in the self-touch condition compared to the misaligned touch condition (*p* = 0.011) (**Figure 2i**). A first control analysis reversed the time axis (**Figure 2j**), after which no positive clusters (self-touch > misaligned touch) were identified, suggesting that the effect is not due to spurious temporal correlations or non-causal dependencies. One negative cluster (misaligned touch > self-touch) was identified, but this did not reject the null hypothesis for the time-reversed Granger causality (*p* = 0.271). A second control analysis reversed the direction of the connectivity from right SI to left cerebellar lobule VI (**Figure 2k**), which found no significant effects (*p* = 0.100), supporting the directional specificity of the main result (from left cerebellum to right SI). Together, these control analyses indicate that the observed directed connectivity is unlikely to be an artefact of temporal structure or of the opposite direction (from right SI to left cerebellum). Directed connectivity plots for stimulus-present and stimulus-absent blocks are shown separately in **Supplementary Figure S2c**, and statistical comparisons are provided in Supplementary Text S3.

### Decreased perceived intensity of self-touch

Finally, participants performed the force-discrimination task that allowed us to quantify their self-touch attenuation (Asimakidou et al., 2022; Bays et al., 2005, 2006; Cemeljic et al., 2025; Job C Kilteni, 2023; Kilteni, 2023; Kilteni et al., 2019, 2020; Kilteni C Ehrsson, 2022; Timar et al., 2023). The task consisted of a two-interval forced-choice task, where participants judged the intensity of two forces, a *test* force of 2 N and a *comparison* force between 1 and 3 N (**Figure 3a**). The task was conducted under three conditions (external touch, self-touch and misaligned touch). A control analysis showed that there were no significant differences in the amount of force used to press with the moving hand during the self-touch and misaligned touch conditions (**Figure 3b**) (*t*(23) = -0.17, *p* = 0.867 *d* = -0.035, *CI^S5^* = [-0.23, 0.20]). Therefore, any perceptual differences detected between the two movement conditions are unlikely to be driven by differences in how participants pressed.

Responses on the force discrimination task (i.e., probability of reporting the second force as stronger than the first as a function of force level) were fitted with psychometric curves (**Figure 3c**), and all fits were considered very good (**Supplementary Figure S3a**). The just noticeable difference (JND) was extracted, which reflects the participants’ discrimination ability. The JNDs did not significantly differ between the conditions (*F*(2, 46) = 0.57, *p* = 0.569, 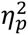 = 0.024) (**Supplementary Figure S3b**).

The point of subjective equality (PSE) was extracted, which represents the intensity at which the *test* force felt as strong as the *comparison* force and quantifies the perceived intensity (i.e. lower PSE values indicate lower perceived intensity). The PSEs significantly differed between the conditions (*F*(2, 46) = 11.95, *p* < 0.001, 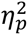 = 0.342). Bonferroni corrected post hoc comparisons revealed significantly lower PSEs in the self-touch condition compared to the external touch condition (*t*(23) = -4.91, *p* < 0.001, *d* = -1.07, *CI^S5^* = [-0.22, -0.07]) (**Figure 3d**). Therefore, in the self-touch condition, participants required a weaker comparison stimulus (compared to the external touch condition) to judge it as equal to the self-touch stimulus, replicating previous behavioural findings using the same force discrimination task (Asimakidou et al., 2022; Bays et al., 2005, 2006; Cemeljic et al., 2025; Job & Kilteni, 2023; Kilteni et al., 2021; Kilteni & Ehrsson, 2022; Timar et al., 2023). The PSEs in the self-touch condition were also significantly lower compared to the misaligned touch condition (*t*(23) = -3.02, *p* = 0.018, *d* = -0.61, *CI^S5^* = [-0.15, -0.01]) (**Figure 3e**), demonstrating that attenuation effects are not due to simultaneous touch on the two hands, temporal predictability, attentional demands, and dual-task requirements. Finally, the PSEs in the external touch condition did not significantly differ from the misaligned touch condition (*t*(23) = 1.94, *p* = 0.194, *d* = 0.46, *CI^S5^* = [-0.02, 0.14]) (**Figure 3f**). Together, the behavioural results show that the touch was attenuated only when the sensorimotor context closely resembled self-touch (i.e., spatially aligned hands touching each other).

## Discussion

We found that self-touch elicited attenuated somatosensory-evoked activity compared to physically identical externally generated touch and spatially misaligned touch. In a separate behavioural task mirroring the MEG conditions, attenuation was similarly reduced when spatial alignment was disrupted. Control analyses did not reveal significant kinematic differences between self-touch and misaligned touch in either task, ruling out motor confounds. Crucially, somatosensory attenuation was *preceded* by reduced pre-stimulus beta power and increased beta-band connectivity from cerebellar lobule VI to primary somatosensory cortex. Given that these effects occurred before touch was received, they suggest that self-touch attenuation is shaped by predictive processes and support a role for the cerebellum in the anticipatory modulation of self-generated somatosensory activity.

The major contribution of our study is the identification of activity that precedes the self-touch input. Beta-band desynchronisation is widely associated with motor preparation and the top-down modulation of sensory processing (Engel & Fries, 2010; Fries, 2015; Saleh et al., 2010). We observed reduced beta power in the self-touch condition relative to the misaligned condition, despite matched motor demands. This suggests that pre-stimulus beta indexes not only motor preparation but also predicted sensory feedback during self-touch. Previous studies have reported reduced beta power before predictable tactile stimuli outside the motor domain (Forster et al., 2025; Kimura, 2021; Kimura & Katayama, 2023; van Ede et al., 2010, 2011b). Our findings extend this work by demonstrating that beta desynchronisation preceding self-touch encodes motor predictions about self-touch, likely modulating expected sensory input. These findings converge on the view that pre-stimulus beta modulations reflect a predictive neural state tailored to expected sensory events.

In addition to beta-band power changes, we found increased pre-stimulus beta-band connectivity between contralateral primary somatosensory cortex and ipsilateral cerebellum in the self-touch vs. misaligned condition. This effect was evident in both coherence and Granger causality, with the latter indicating directed connectivity from the left cerebellar lobule VI to the contralateral primary somatosensory cortex. Directed connectivity measures like Granger causality can be sensitive to shared temporal structure, so we ran two control analyses (time reversal and direction reversal), neither of which produced a similar result, supporting the specificity of the finding. We interpret this directed connectivity as complementary to the pre-stimulus coherence beta effects, adding mechanistic directionality. These findings support the forward model framework, which posits that the cerebellum generates predictive signals to modulate the somatosensory cortex in anticipation of self-generated sensory input. Unlike previous neuroimaging work limited to post-stimulus activity (Hesse et al., 2010), or averaged over pre- and post-stimulus activity (Blakemore et al., 1998; Kilteni & Ehrsson, 2020), we show that cerebellar-somatosensory connectivity occurs *before* the sensory event and is *directed* from the cerebellum to the somatosensory cortex, two criteria important for establishing the predictive nature of the effect. One plausible anatomical route is via cerebello–thalamo–cortical pathways that convey cerebellar output to sensorimotor regions (Andersen & Dalal, 2024; Aoki et al., 2019; Bostan et al., 2013; Dum & Strick, 2003; Keser et al., 2015), which could support rapid pre-stimulus influences on SI. Additional analyses of the separate stimulus-present and stimulus-absent conditions suggest that the self-touch vs misaligned touch coherence effect may be partly driven by lower coherence in the misaligned stimulus-present condition relative to its stimulus-absent control, rather than solely by increased coherence during self-touch. This may indicate that spatially incongruent action-touch mappings alter ongoing cerebellar-somatosensory coupling in a way that is not captured by a simple increase in connectivity during self-touch. By contrast, the Granger causality comparisons were more consistent with increased directed connectivity from the cerebellum to SI during self-touch, supporting the interpretation that predictive cerebellar influences are enhanced before spatially congruent self-touch.

Behaviourally, participants perceived self-touch as weaker than externally generated touch, replicating previous findings (Asimakidou et al., 2022; Bays et al., 2005, 2006; Cemeljic et al., 2025; Kilteni et al., 2018, 2019, 2021; Kilteni & Ehrsson, 2017, 2020, 2022). Attenuation was reduced when the hands were misaligned, indicating that attenuation depends on the spatial correspondence between movement and tactile feedback. In parallel, early somatosensory-evoked activity was attenuated for self-touch relative to external touch, consistent with prior neuroimaging findings (Hesse et al., 2010). This early attenuation was significantly diminished in the misaligned condition, supporting the interpretation that it reflects the precision of action-based predictions rather than non-predictive differences between self-touch and external touch. Many alternative explanations that differ between self-touch and external touch are held constant in the misaligned condition, including movement, bilateral tactile stimulation, temporal predictability, divided attention, agency, and general expectation of touch. Notably, no significant difference was observed between self-touch and misaligned touch at the later M100 component, suggesting that spatial specificity primarily affects early cortical somatosensory processing. Together, the behavioural and neural findings reinforce the idea that attenuation depends on the correspondence between predicted and actual sensory feedback (Bays & Wolpert, 2008; Blakemore et al., 2000; Wolpert & Flanagan, 2001).

Predictions about the sensory consequences of our own movements are shaped by long-term experience of the typical relationships between movements and sensations. However, rapidly learned contingencies within a particular task can also affect perception. Behavioural work on temporal delays in self-touch shows that repeated exposure to a delay can partly reduce attenuation for immediate touch and increase attenuation for delayed touch, with the magnitude of recalibration depending on the number of exposure trials (Kilteni et al., 2019; Kilteni & Ehrsson, 2024). We manipulated the spatial relationship between the moving hand and the stimulated site while keeping the temporal contingency constant. Participants could, in principle, learn to attenuate the misaligned touch to some extent. Nevertheless, this hypothesized recalibration would require repeated exposure to such trials over a prolonged period (e.g., >400 trials in Kilteni et al., 2019), whereas the brief presentation of trials used here would not be sufficient. This is supported by our behavioural results and early evoked neural responses, which showed significant differences between the self-touch and misaligned conditions despite exposure to misaligned trials. Importantly, the misaligned mapping continued to violate deeply ingrained spatial regularities between hand movements and touch at the body location where contact occurs. Lifelong experience of spatially aligned touch is likely to provide the forward model with strong body-centred priors that action-effects occur at the effector’s contact site. We acknowledge that this spatial manipulation provides an indirect manipulation of prediction strength. Accordingly, rather than implying that misaligned touch is entirely “unpredicted” while self-touch is “predicted”, our findings indicate relative differences in the strength and precision of spatial action-based predictions across conditions that share a general expectation that a touch will occur.

Our findings align with animal studies showing reduced neural responses to self-generated stimuli across sensory modalities, including vestibular, auditory, tactile, mechanosensory and electrosensory domains (Audette et al., 2022; Brooks et al., 2015; Kilteni et al., 2025; Perks et al., 2020; Schneider et al., 2018; Singla et al., 2017). Studies have also identified neural signals that precede and predict expected sensory consequences of action, such as predictive representations of self-generated sounds in the auditory cortex (Audette et al., 2022). Previous studies also implicate cerebellar or cerebellar-like structures in predictive sensory attenuation, including suppression of vestibular input during head movements (Brooks et al., 2015) and cancellation of self-generated sounds or electrosensory signals by cerebellum-like circuits (Singla et al., 2017; Wallach & Sawtell, 2023). Together with our observation of increased directed connectivity from the cerebellum to the primary somatosensory cortex, this suggests that a predictive cerebellar modulation of early sensory processing may be a conserved feature across species and sensory modalities.

Disruptions in the integration of predictive signals with sensory input have been central to neurobiological accounts of psychosis spectrum disorders, including schizophrenia (Blakemore et al., 2000; Corlett et al., 2019; Shergill et al., 2005, 2014) and schizotypy (Asimakidou et al., 2022). Reduced attenuation of self-generated auditory and somatosensory stimuli has been reported in schizophrenia (Shergill et al., 2014; Whitford et al., 2011), consistent with proposals that impaired prediction contributes to altered self-other distinction, weakened agency, and symptoms such as hallucinations or delusions of control (Frith, 2005a, 2005b; Leptourgos & Corlett, 2020; Poletti et al., 2019). Most previous neuroimaging studies have focused on post-stimulus activity, leaving open whether earlier predictive processes are compromised. Future studies should investigate whether pre-stimulus beta desynchronization and cerebellar-somatosensory connectivity are diminished or absent in individuals with schizophrenia or elevated schizotypal traits, potentially offering early indicators of disrupted prediction in clinical populations.

In summary, self-touch was perceived as less intense, evoked weaker somatosensory activity, and was preceded by beta-band desynchronisation and enhanced directed cerebellar-to-somatosensory connectivity. These effects were reduced when the spatial alignment between movement and touch was disrupted, indicating dependence on precise motor predictions. By isolating pre-stimulus activity and providing evidence for cerebellar-to-cortical information flow, our results extend previous work and provide a mechanism by which the brain anticipates self-generated tactile events. Such predictive processes may be fundamental to distinguishing self-from externally generated events, and the pre-stimulus markers identified here offer promise for understanding disrupted sensory predictions in psychosis spectrum disorders.

**Conflict of interest statement**

The authors declare no competing financial interests

## Acknowledgements

We thank Lili Timar for her assistance with collecting the behavioural data. Experimental costs were covered by the Swedish Research Council (VR 2019-01909 and VR 2024-00906). X.J. was supported by the Marie Skłodowska-Curie Intra-European Individual Fellowship (grant number 101059348) and The Strategic Research Area Neuroscience (StratNeuro). K.K was supported by the European Research Council (ERC 101039152).

## References

Andersen, L. M., & Dalal, S. S. (2021). The cerebellar clock: Predicting and timing somatosensory touch. NeuroImage, 238. 10.1016/j.neuroimage.2021.118202

Andersen, L. M., & Dalal, S. S. (2024a). Detection of Threshold-Level Stimuli Modulated by Temporal Predictions of the Cerebellum. ENeuro, 11(4). 10.1523/ENEURO.0070-24.2024

Andersen, L. M., & Dalal, S. S. (2024b). The role of the cerebellum in timing. Current Opinion in Behavioral Sciences, *5S*, 101427. 10.1016/j.cobeha.2024.101427

Andersen, L. M., Jerbi, K., & Dalal, S. S. (2020). Can EEG and MEG detect signals from the human cerebellum? NeuroImage, 215, 116817. 10.1016/j.neuroimage.2020.116817

Andersen, L. M., & Lundqvist, D. (2019). Somatosensory responses to nothing: An MEG study of expectations during omission of tactile stimulations. NeuroImage, 184, 78–89. 10.1016/j.neuroimage.2018.09.014

Aoki, S., Coulon, P., & Ruigrok, T. J. H. (2019). Multizonal Cerebellar Influence Over Sensorimotor Areas of the Rat Cerebral Cortex. Cerebral Cortex, *2S*(2). 598–614. 10.1093/cercor/bhx343

Asimakidou, E., Job, X., & Kilteni, K. (2022). The positive dimension of schizotypy is associated with a reduced attenuation and precision of self-generated touch. Schizophrenia, 8(1), 57. 10.1038/s41537-022-00264-6

Audette, N. J., Zhou, W. X., La Chioma, A., & Schneider, D. M. (2022). Precise movement-based predictions in the mouse auditory cortex. Current Biology, 32(22), 4925–4940. 10.1016/j.cub.2022.09.064

Bays, P. M., Flanagan, J. R., & Wolpert, D. M. (2006). Attenuation of self-generated tactile sensations is predictive, not postdictive. PLoS Biology, 4(2), e28. 10.1371/journal.pbio.0040028

Bays, P. M., & Wolpert, D. M. (2008). Predictive attenuation in the perception of touch. Sensorimotor Foundations of Higher Cognition, 22, 339–358. 10.1093/acprof:oso/9780199231447.003.0016

Bays, P. M., Wolpert, D. M., & Flanagan, J. R. (2005). Perception of the consequences of self-action is temporally tuned and event driven. Current Biology, 15(12), 1125–1128. 10.1016/j.cub.2005.05.023

Blakemore, S. J., Frith, C. D., & Wolpert, D. M. (1999). Spatio-temporal prediction modulates the perception of self-produced stimuli. Journal of Cognitive Neuroscience, 11(5), 551–559. 10.1162/089892999563607

Blakemore, S. J., Wolpert, D., & Frith, C. (2000). Why can’t you tickle yourself? NeuroReport, 11(11). R11–R16. 10.1097/00001756-200008030-00002

Blakemore, S. J., Wolpert, D. M., & Frith, C. D. (1998). Central cancellation of self-produced tickle sensation. Nature Neuroscience, 1(7), 635–640. 10.1038/2870

Blakemore, S. J., Smith, J., Steel, R., Johnstone, E. C., & Frith, C. D. (2000). The perception of self-produced sensory stimuli in patients with auditory hallucinations and passivity experiences: evidence for a breakdown in self-monitoring. Psychological Medicine, 30(5), 1131–1139.

Bostan, A. C., Dum, R. P., & Strick, P. L. (2013). Cerebellar networks with the cerebral cortex and basal ganglia. Trends in Cognitive Sciences, 17(5), 241–254. 10.1016/j.tics.2013.03.003

Brooks, J. X., Carriot, J., & Cullen, K. E. (2015). Learning to expect the unexpected: rapid updating in primate cerebellum during voluntary self-motion. Nature Neuroscience, 18(9), 1310–1317. 10.1038/nn.4077

Cemeljic, N., Job, X., & Kilteni, K. (2025). Predictions of bimanual self-touch determine the temporal tuning of somatosensory perception. IScience, 28(2), 111643. 10.1016/j.isci.2024.111643

Corlett, P. R., Horga, G., Fletcher, P. C., Alderson-Day, B., Schmack, K., & Powers, A. R. (2019). Hallucinations and Strong Priors. Trends in Cognitive Sciences, 23(2), 114–127. 10.1016/j.tics.2018.12.001

Dum, R. P., & Strick, P. L. (2003). An unfolded map of the cerebellar dentate nucleus and its projections to the cerebral cortex. Journal of Neurophysiology, *8S*(1), 634-639. 10.1152/jn.00626.2002

Engel, A. K., & Fries, P. (2010). Beta-band oscillations-signalling the status quo? In Current Opinion in Neurobiology, 20(2), 156–165. 10.1016/j.conb.2010.02.015

Engel, A. K., Fries, P., & Singer, W. (2001). Dynamic predictions: Oscillations and synchrony in top–down processing. Nature Reviews Neuroscience, 2(10). 10.1038/35094565

Fletcher, P. C., & Frith, C. D. (2009). Perceiving is believing: A Bayesian approach to explaining the positive symptoms of schizophrenia. Nature Reviews Neuroscience, 10(1), 704–716. 10.1038/nrn2536

Forster, C., Stephani, T., Grund, M., Panagoulas, E., Al, E., Hofmann, S. M., Nikulin, V. V., & Villringer, A. (2025). Pre-stimulus beta power mediates explicit and implicit perceptual biases in distinct cortical areas. Communications Psychology, 3(1), 93. 10.1038/s44271-025-00265-y

Franklin, D. W., & Wolpert, D. M. (2011). Computational mechanisms of sensorimotor control. Neuron, 72(3), 425–442. 10.1016/j.neuron.2011.10.006

Fries, P. (2015). Rhythms for Cognition: Communication through Coherence. Neuron, 88(1), 220–235. 10.1016/j.neuron.2015.09.034

Frith, C. (2005a). The neural basis of hallucinations and delusions. Comptes Rendus Biologies, 328(2), 169–175. 10.1016/j.crvi.2004.10.012

Frith, C. (2005b). The self in action: Lessons from delusions of control. Consciousness and Cognition, 14(4), 752–770. 10.1016/j.concog.2005.04.002

Frith, C. D. (2019). Can a problem with corollary discharge explain the symptoms of schizophrenia? Biological Psychiatry: Cognitive Neuroscience and Neuroimaging, 4(9), 768–769. 10.1016/j.bpsc.2019.07.003

Frith, C. D., Blakemore, S. J., & Wolpert, D. M. (2000). Explaining the symptoms of schizophrenia: Abnormalities in the awareness of action. Brain Research Reviews, 31(2–3), 357–363. 10.1016/S0165-0173(99)00052-1

Grodd, W., Hülsmann, E., Lotze, M., Wildgruber, D., & Erb, M. (2001). Sensorimotor mapping of the human cerebellum: fMRI evidence of somatotopic organization. Human Brain Mapping, 13(2), 55–73. 10.1002/hbm.1025

Gross, J., Kujala, J., Hämäläinen, M., Timmermann, L., Schnitzler, A., & Salmelin, R. (2001). Dynamic imaging of coherent sources: Studying neural interactions in the human brain. *Proceedings of the National Academy of Sciences of the United States of America*, S8(2), 694–699. 10.1073/pnas.98.2.694

Gupta, K., Chowdhury, R. R., Chakrabarti, S., & Schwarz, C. (2023). Discerning state estimation and sensory gating, two presumptive predictive signals in mouse barrel cortex. BioRxiv, 2023.12.23.573180. 10.1101/2023.12.23.573180

Hesse, M. D., Nishitani, N., Fink, G. R., Jousmäki, V., & Hari, R. (2010). Attenuation of somatosensory responses to self-produced tactile stimulation. Cerebral Cortex, 20(2), 425–432. 10.1093/cercor/bhp110

Job, X., & Kilteni, K. (2023). Action does not enhance but attenuates predicted touch. ELife, 12, e90912. 10.7554/eLife.90912

Kalcher, J., & Pfurtscheller, G. (1995). Discrimination between phase-locked and non-phase-locked event-related EEG activity. *Electroencephalography and Clinical Neurophysiology*, 94(5), 381–384. 10.1016/0013-4694(95)00040-6

Keser, Z., Hasan, K. M., Mwangi, B. I., Kamali, A., Ucisik-Keser, F. E., Riascos, R. F., Yozbatiran, N., Francisco, G. E., & Narayana, P. A. (2015). Diffusion tensor imaging of the human cerebellar pathways and their interplay with cerebral macrostructure. *Frontiers in Neuroanatomy*, S, 41. 10.3389/fnana.2015.00041

Kilteni, K. (2023). Methods of Somatosensory Attenuation. In Somatosensory Research Methods (pp. 35–53). New York, NY: Springer US. 10.1007/978-1-0716-3068-6_2

Kilteni, K. (2025). The extraordinary enigma of ordinary tickle behavior: Why gargalesis still puzzles neuroscience. Science Advances, 11(21), eadt0350. 10.1126/sciadv.adt0350

Kilteni, K., Andersson, B. J., Houborg, C., & Ehrsson, H. H. (2018). Motor imagery involves predicting the sensory consequences of the imagined movement. *Nature Communications*, S(1), 1617. 10.1038/s41467-018-03989-0

Kilteni, K., & Ehrsson, H. H. (2017). Sensorimotor predictions and tool use: Hand-held tools attenuate self-touch. Cognition, *1C5*, 1–9. 10.1016/j.cognition.2017.04.005

Kilteni, K., & Ehrsson, H. H. (2020). Functional Connectivity between the Cerebellum and Somatosensory Areas Implements the Attenuation of Self-Generated Touch. The Journal of Neuroscience, 40(4), 894–906. 10.1523/JNEUROSCI.1732-19.2019

Kilteni, K., & Ehrsson, H. H. (2022). Predictive attenuation of touch and tactile gating are distinct perceptual phenomena. IScience, 25(4), 104077. 10.1016/j.isci.2022.104077

Kilteni, K., & Ehrsson, H. H. (2024). Dynamic changes in somatosensory and cerebellar activity mediate temporal recalibration of self-touch. Communications Biology, 7(1), 522. 10.1038/s42003-024-06188-4

Kilteni, K., Engeler, P., Boberg, I., Maurex, L., & Ehrsson, H. H. (2021). No evidence for somatosensory attenuation during action observation of self-touch. European Journal of Neuroscience, 54(7), 6422–6444. 10.1111/ejn.15436

Kilteni, K., Engeler, P., & Ehrsson, H. H. (2020). Efference Copy Is Necessary for the Attenuation of Self-Generated Touch. IScience, 23(2), 100843. 10.1016/j.isci.2020.100843

Kilteni, K., Houborg, C., & Ehrsson, H. H. (2019). Rapid learning and unlearning of predicted sensory delays in self-generated touch. ELife, 8, e42888. 10.7554/eLife.42888

Kilteni, K., Houborg, C., & Ehrsson, H. H. (2023). Brief temporal perturbations in somatosensory reafference disrupt perceptual and neural attenuation and increase supplementary motor area–cerebellar connectivity. The Journal of Neuroscience, 43(28), 5251–5263 10.1523/jneurosci.1743-22.2023

Kimura, T. (2021). Approach of visual stimuli facilitates the prediction of tactile events and suppresses beta band oscillations around the primary somatosensory area. NeuroReport, 32(7), 631–635. 10.1097/WNR.0000000000001643

Kimura, T., & Katayama, J. (2023). Visual stimuli in the peripersonal space facilitate the spatial prediction of tactile events—A comparison between approach and nearness effects. Frontiers in Human Neuroscience, 17, 1203100. 10.3389/fnhum.2023.1203100

Leptourgos, P., & Corlett, P. R. (2020). Embodied predictions, agency, and psychosis. Frontiers in Big Data, 3, 27. 10.3389/fdata.2020.00027

Manni, E., & Petrosini, L. (2004). A century of cerebellar somatotopy: A debated representation. Nature Reviews Neuroscience, 5(3) 241–249. 10.1038/nrn1347

Maris, E., & Oostenveld, R. (2007). Nonparametric statistical testing of EEG- and MEG-data. Journal of Neuroscience Methods, *1C4*(1), 177–190. 10.1016/j.jneumeth.2007.03.024

McNamee, D., & Wolpert, D. M. (2019). Internal models in biological control. *Annual Review of Control*, Robotics, and Autonomous Systems, 2(1), 339–364. 10.1146/annurev-control-060117-105206

Miall, R. C., & Wolpert, D. M. (1996). Forward models for physiological motor control. *Neural Networks*, S(8), 1265–1279. 10.1016/S0893-6080(96)00035-4

Mottolese, C., Richard, N., Harquel, S., Szathmari, A., Sirigu, A., & Desmurget, M. (2013). Mapping motor representations in the human cerebellum. Brain, *13C*(1), 330–342. 10.1093/brain/aws186

Nolte, G. (2003). The magnetic lead field theorem in the quasi-static approximation and its use for magnetoenchephalography forward calculation in realistic volume conductors. Physics in Medicine and Biology, 48(22), 3637. 10.1088/0031-9155/48/22/002

Oldfield, R. C. (1971). The assessment and analysis of handedness: The Edinburgh inventory. *Neuropsychologia*, S(1), 97–113. 10.1016/0028-3932(71)90067-4

Oostenveld, R., Fries, P., Maris, E., & Schoffelen, J. M. (2011). FieldTrip: Open source software for advanced analysis of MEG, EEG, and invasive electrophysiological data. Computational Intelligence and Neuroscience, 2011(1), 156869. 10.1155/2011/156869

Pando-Naude, V., & Andersen, L. M. (2025). Somatosensory rhythms and cerebellar-basal ganglia beta-band interactions in Parkinson’s disease, bioRxiv, 2025-05. 10.1101/2025.05.15.653735

Perks, K. E., Krotinger, A., & Bodznick, D. (2020). A cerebellum-like circuit in the lateral line system of fish cancels mechanosensory input associated with its own movements. Journal of Experimental Biology, 223(4), jeb204438. 10.1242/jeb.204438

Pfurtscheller, G., & Lopes Da Silva, F. H. (1999). Event-related EEG/MEG synchronization and desynchronization: Basic principles. Clinical Neurophysiology, 110(11), 1842–1857. 10.1016/S1388-2457(99)00141-8

Poletti, M., Tortorella, A., & Raballo, A. (2019). Impaired Corollary Discharge in Psychosis and At-Risk States: Integrating Neurodevelopmental, Phenomenological, and Clinical Perspectives. Biological Psychiatry: Cognitive Neuroscience and Neuroimaging, 4(9), 832–841. 10.1016/j.bpsc.2019.05.008

Quintana, D. S., & Williams, D. R. (2018). Bayesian alternatives for common null-hypothesis significance tests in psychiatry: A non-technical guide using JASP. BMC Psychiatry, 18(1), 178. 10.1186/s12888-018-1761-4

Raine, A. (1991). The spq: A scale for the assessment of schizotypal personality based on DSM-III-r criteria. Schizophrenia Bulletin, 17(4), 555–564. 10.1093/schbul/17.4.555

Saleh, M., Reimer, J., Penn, R., Ojakangas, C. L., & Hatsopoulos, N. G. (2010). Fast and Slow Oscillations in Human Primary Motor Cortex Predict Oncoming Behaviorally Relevant Cues. *Neuron*, C5(4), 461–471. 10.1016/j.neuron.2010.02.001

Samuelsson, J. G., Sundaram, P., Khan, S., Sereno, M. I., & Hämäläinen, M. S. (2020). Detectability of cerebellar activity with magnetoencephalography and electroencephalography. Human Brain Mapping, 41(9), 2357–2372. 10.1002/hbm.24951

Schneider, D. M., Sundararajan, J., & Mooney, R. (2018). A cortical filter that learns to suppress the acoustic consequences of movement. Nature, 5C1(7723), 391–395. 10.1038/s41586-018-0520-5

Shergill, S. S., Bays, P. M., Frith, C. D., & Wolpert, D. M. (2003). Two eyes for an eye: the neuroscience of force escalation. Science 301(5630), 187. 10.1126/science.1085327

Shergill, S. S., Samson, G., Bays, P. M., Frith, C. D., & Wolpert, D. M. (2005). Evidence for sensory prediction deficits in schizophrenia. American Journal of Psychiatry, *1C2*(12), 2384–2386. 10.1176/appi.ajp.162.12.2384

Shergill, S. S., White, T. P., Joyce, D. W., Bays, P. M., Wolpert, D. M., & Frith, C. D. (2014). Functional magnetic resonance imaging of impaired sensory prediction in schizophrenia. JAMA Psychiatry, 71(1), 28–35. 10.1001/jamapsychiatry.2013.2974

Singla, S., Dempsey, C., Warren, R., Enikolopov, A. G., & Sawtell, N. B. (2017). A cerebellum-like circuit in the auditory system cancels responses to self-generated sounds. Nature Neuroscience, 20(7), 943–950. 10.1038/nn.4567

Spitzer, B., & Haegens, S. (2017). Beyond the status quo: A role for beta oscillations in endogenous content (RE)activation. eNeuro, 4(4). 10.1523/ENEURO.0170-17.2017

Taulu, S., Kajola, M., & Simola, J. (2004). Suppression of interference and artifacts by the signal space separation method. Brain Topography, *1C*(4), 269–275. 10.1023/B:BRAT.0000032864.93890.f9

Taulu, S., & Simola, J. (2006). Spatiotemporal signal space separation method for rejecting nearby interference in MEG measurements. Physics in Medicine and Biology, 51(7), 1759. 10.1088/0031-9155/51/7/008

Timar, L., Job, X., Orban de Xivry, J.-J., & Kilteni, K. (2023). Aging exerts a limited influence on the perception of self-generated and externally generated touch. Journal of Neurophysiology, 130(4), 871–882. 10.1152/jn.00145.2023

Valè, nicola, Tomic, I., Gironés, Z., Wolpert, D. M., Kilteni, K., & Bays, P. M. (2025). Divisive attenuation based on noisy sensorimotor predictions accounts for excess variability in self-touch. Journal of Neurophysiology, 130(4), 871–882. 10.1152/jn.00055.2025

van Doorn, J., van den Bergh, D., Böhm, U., Dablander, F., Derks, K., Draws, T., Etz, A., Evans, N. J., Gronau, Q. F., Haaf, J. M., Hinne, M., Kucharský, Š., Ly, A., Marsman, M., Matzke, D., Gupta, A. R. K. N., Sarafoglou, A., Stefan, A., Voelkel, J. G., & Wagenmakers, E. J. (2021). The JASP guidelines for conducting and reporting a Bayesian analysis. Psychonomic Bulletin and Review, 28(3), 813–826. 10.3758/s13423-020-01798-5

van Ede, F., de Lange, F. P., Jensen, O., & Maris, E. (2011a). Orienting attention to an upcoming tactile event involves a spatially and temporally specific modulation of sensorimotor alpha- and beta-band oscillations. Journal of Neuroscience, 31(6), 2016–2024. 10.1523/JNEUROSCI.5630-10.2011

van Ede, F., de Lange, F. P., Jensen, O., & Maris, E. (2011b). Orienting attention to an upcoming tactile event involves a spatially and temporally specific modulation of sensorimotor alpha- and beta-band oscillations. Journal of Neuroscience, 31(6), 2016–2024. 10.1523/JNEUROSCI.5630-10.2011

van Ede, F., Jensen, O., & Maris, E. (2010). Tactile expectation modulates pre-stimulus Α-band oscillations in human sensorimotor cortex. NeuroImage, 51(2) 867–876. 10.1016/j.neuroimage.2010.02.053

Van Veen, B. D., Van Drongelen, W., Yuchtman, M., & Suzuki, A. (1997). Localization of brain electrical activity via linearly constrained minimum variance spatial filtering. IEEE Transactions on Biomedical Engineering, 44(9), 867–880. 10.1109/10.623056

Wallach, A., & Sawtell, N. B. (2023). An internal model for canceling self-generated sensory input in freely behaving electric fish. Neuron, 111(16), 2570–2582. 10.1016/j.neuron.2023.05.019

Weiskrantz, L., Elliott, J., & Darlington, C. (1971). Preliminary observations on tickling oneself. Nature, 230(5296), 598–599. 10.1038/230598a0

Whitford, T. J., Mathalon, D. H., Shenton, M. E., Roach, B. J., Bammer, R., Adcock, R. A., Bouix, S., Kubicki, M., De Siebenthal, J., Rausch, A. C., Schneiderman, J. S., & Ford, J. M. (2011). Electrophysiological and diffusion tensor imaging evidence of delayed corollary discharges in patients with schizophrenia. Psychological Medicine, 41(5), 959–969. 10.1017/S0033291710001376

Wolpert, D. M., & Flanagan, J. R. (2001). Motor prediction. Current Biology, 11(18), R729–R732. 10.1016/S0960-9822(01)00432-8

Yamashiro, K., Sato, D., Onishi, H., Sugawara, K., Otsuru, N., Kirimoto, H., Nakazawa, S., Yamazaki, Y., Shirozu, H., & Maruyama, A. (2019). Change-driven M100 component in the bilateral secondary somatosensory cortex: A magnetoencephalography study. Brain Topography, 32(2), 435–444. 10.1007/s10548-018-0687-y

