## Supplementary material for "Motor prediction reduces beta-band power and enhances cerebellar-somatosensory connectivity before self-touch to enable its attenuation"

**Supplementary Text S1**. Control analyses for movement during the MEG session.

*Movement onset and maximum acceleration*

An accelerometer, attached to the dorsal surface of the middle phalanx of the right index finger, measured the acceleration of the finger movements. Movement onset was calculated as the first time point that acceleration exceeded 10% of the total amplitude for a duration of 100 ms. **Figure S1a** (upper panel) shows the average onset and maximum acceleration of the moving hand for the conditions involving movement, as well as the average acceleration across time (lower panel).

A repeated measures analysis of variance (rmANOVA) for the movement onset did not reveal a significant main effect of condition ($F(1, 23) = 0.90$, $p = 0.353$, $\eta_p^2 = 0.038$), or a main effect of stimulus presence ($F(1, 23) = 0.21$, $p = 0.654$, $\eta_p^2 = 0.009$), or an interaction between condition and stimulus presence ($F(1, 23) = 0.38$, $p = 0.542$, $\eta_p^2 = 0.016$). This suggests that participants did not take longer to initiate their movement in any of the conditions or in trials with/without the tactile stimulus.

A rmANOVA for the maximum acceleration did not yield a significant main effect of condition ($F(1, 23) = 0.43$, $p = 0.521$, $\eta_p^2 = 0.018$), or a main effect of stimulus presence ($F(1, 23) = 0.38$, $p = 0.544$, $\eta_p^2 = 0.016$), or an interaction between condition and stimulus presence ($F(1, 23) = 0.01$, $p = 0.929$, $\eta_p^2 = 0.001$). This suggests participants did not accelerate faster in either of the movement conditions or in trials with/without the tactile stimulus.

*Movement intensity*

An optical fiber measured the amplitude of the active finger press (i.e., how forceful the participant pressed on the custom lever device) as a function of time relative to triggering the response device. Note that the amplitude of the active (right) finger press was not linked to the magnitude of the tactile stimulus delivered to the passive (left) finger, which was held constant. Participants were trained before the experiment regarding how to press the lever and were asked to be consistent across the experiment. If the participants pressed less than requested during the experiment, they were given feedback to correct their movement. The magnitude of the touch applied on the left index finger was kept constant to permit the analysis of the brain responses. **Figure S1b** (upper panel) shows the average maximum amplitude of the active finger press for the conditions involving movement and the average amplitude across time (lower panel). Triggering the response device (i.e., 0 seconds) either caused a tactile stimulus on the left index finger (stimulus-present) or not (stimulus-absent). We compared the conditions with action (self-touch vs. misaligned touch) for stimulus-present and stimulus-absent blocks.

A rm ANOVA on the maximum amplitudes did not yield a significant main effect of condition ($F(1, 23) = 0.97$, $p = 0.335$, $\eta_p^2 = 0.040$). There was a significant main effect of stimulus presence with larger amplitude presses in the stimulus-absent blocks compared to the stimulus-present blocks ($F(1, 23) = 12.37$, $p = 0.002$, $\eta_p^2 = 0.350$). However, there was no significant interaction ($F(1, 23) = 2.05$, $p = 0.165$, $\eta_p^2 = 0.082$). Although we observed a main effect of stimulus presence, indicating generally larger amplitude presses in the stimulus-absent blocks compared to stimulus present blocks, this effect did not interact with condition. Therefore, the main effect of stimulus presence cannot account for the condition-specific MEG effects observed in our analyses.

*Inter-stimulus-interval*

The mean inter-stimulus interval for the self-touch stimulus-present blocks was 2692 ms ± 149 SD, the self-touch stimulus-absent blocks was 2688 ms ± 156 SD, the misaligned touch stimulus-present blocks was 2701 ms ± 128 SD, and the misaligned touch stimulus-absent blocks was 2711 ms ± 128 SD. A rmANOVA comparing the inter-stimulus intervals with a factor of condition (self-touch vs. misaligned touch) and stimulus presence (stimulus-present vs. stimulus-absent) showed no significant main effect of condition ($F(1, 23) = 0.89$, $p = 0.355$, $\eta_p^2 = 0.037$), or stimulus presence ($F(1, 23) = 0.08$, $p = 0.784$, $\eta_p^2 = 0.003$) and no significant interaction ($F(1, 23) = 1.67$, $p = 0.209$, $\eta_p^2 = 0.068$).

**Supplementary Text S2.** Additional analyses of pre-stimulus time-frequency responses.

We note that comparing pre-stimulus activity between a condition with movement (e.g., self-touch condition) and the condition without any movement (external touch condition) is difficult to interpret because any pre-stimulus differences observed may reflect an unknown combination of movement-related activity and predictive attenuation of the touch. Furthermore, the external touch condition had a fixed cue-onset interval of 500 ms while the movement conditions had a variable cue-onset interval, given that participants had to trigger the touch by pressing the lever, the interval depended on their movement time. Pre-stimulus differences are therefore confounded by differences in the cue-locked activity. Despite these confounds, for completeness, we also compared pre-stimulus beta-band (13-30 Hz) activity in the external touch condition and movement conditions (self-touch and misaligned touch conditions). Interestingly, for the comparison between self-touch and external touch conditions, a cluster-based permutation test showed a significant difference between the conditions with greater beta desynchrony in the self-touch condition ($p = 0.007$). This cluster resembled that of the comparison between the self-touch and the misaligned touch condition. For the comparison between the misaligned touch and external touch conditions, a cluster-based permutation test showed significantly greater beta synchrony in the misaligned touch condition compared to the self-touch condition ($p = 0.022$).

**Supplementary Text S3.** Additional analyses of stimulus-present and stimulus-absent connectivity

We examined whether coherence and Granger causality differed between the self-touch and misaligned-touch conditions separately for stimulus-present and stimulus-absent trials, and whether stimulus-present trials differed from stimulus-absent trials within each condition.
For the direct comparison between self-touch and misaligned-touch stimulus-present trials, cluster permutation tests indicated marginal but non-significant effects for both coherence ($p$ = 0.060) and Granger causality ($p$ = 0.091). The corresponding comparisons between self-touch and misaligned-touch stimulus-absent trials were non-significant for both coherence (all $p$ > 0.154) and Granger causality (all $p$ > 0.100).

We then compared stimulus-present and stimulus-absent trials within each movement condition. For coherence, the self-touch condition did not show a significant difference between stimulus-present and stimulus-absent trials ($p$ = 0.214), whereas the misaligned-touch condition showed significantly lower coherence for stimulus-present than stimulus-absent trials ($p$ = 0.017). For Granger causality, the self-touch condition showed significantly higher values for stimulus-present than stimulus-absent trials ($p$ = 0.044), whereas the corresponding comparison in the misaligned-touch condition was not significant (all $p$ > 0.127).

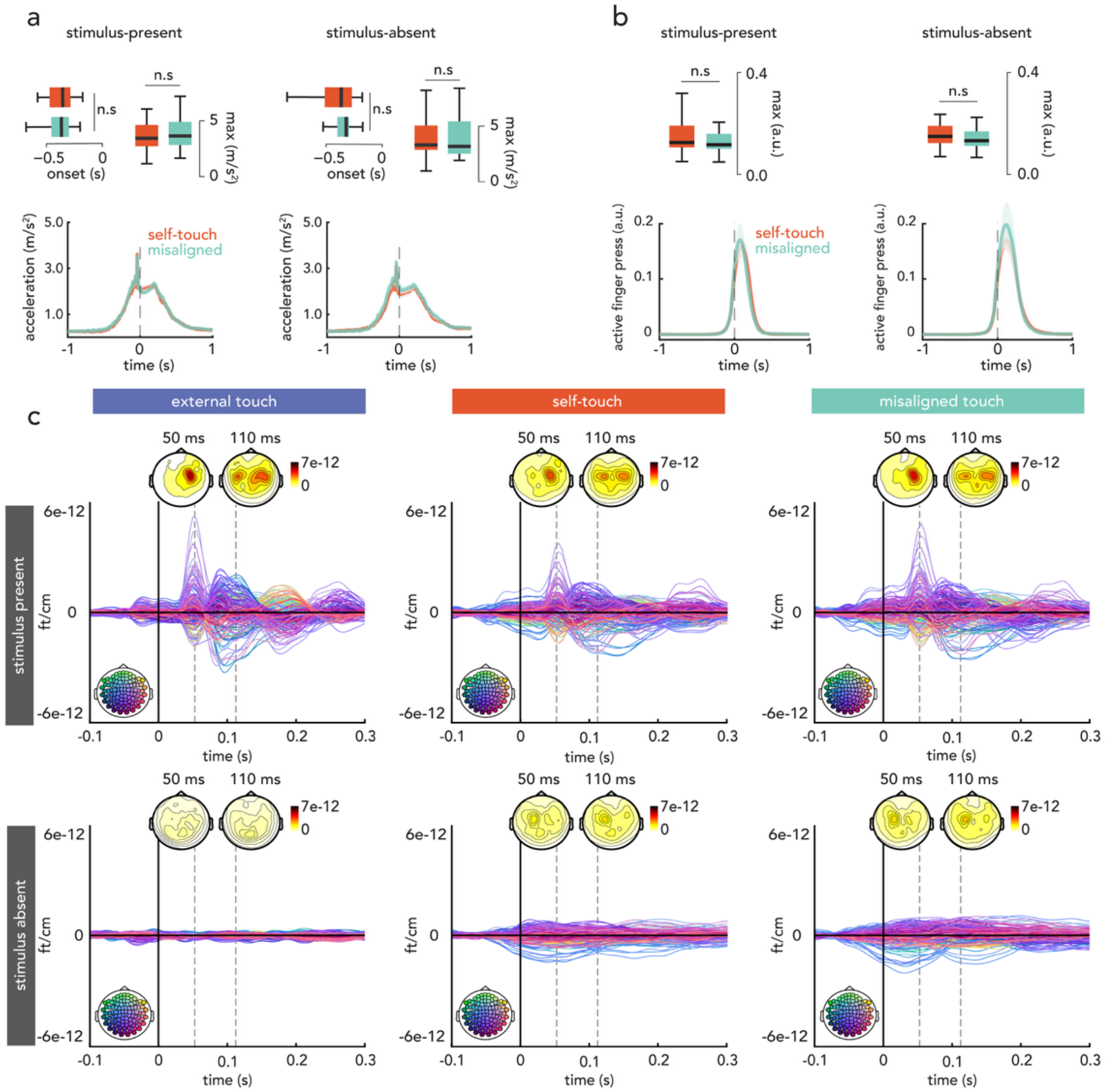


**Supplementary Figure S1. Movement kinematics and event-related fields. (a)** Acceleration of the moving right index finger across time in the movement conditions (self-touch and misaligned touch). The dashed line (0 seconds) is the timepoint at which the right index finger pressed the response device to either trigger a touch (stimulus-present) or not (stimulus-absent). Box plots show the median and interquartile range for the movement onset (i.e. the first time point that acceleration exceeded 10% of the total amplitude for a duration of 100 ms) and the maximum acceleration. **(b)** Optical fiber traces from the response device lever measure the amplitude of the active finger press (i.e., how forceful the participant pressed on the lever device) as a function of time relative to triggering the response device. Box plots show the median and interquartile range for the maximum amplitude. **(a, b)** Shaded error bands show ± standard error of the mean (s.e.m.). **(c)** Butterfly plots of the 204 gradiometers. Upper panel: Stimulus-present blocks: time-locked to the onset of the tactile stimulus (dashed line) on the left index finger with topographical plots of the root mean square values of the combined gradiometer pairs. Lower panel: Stimulus-absent blocks: time-locked to the moment the touch would have been delivered in the stimulus-present blocks. Topographies show the M50 (50 ms) and M100 (110 ms) components of the event-related fields. Coloured heads in the bottom-left corners indicate sensor locations. In the main analysis, stimulus-absent waveforms were subtracted from stimulus-present waveforms before comparing the conditions.

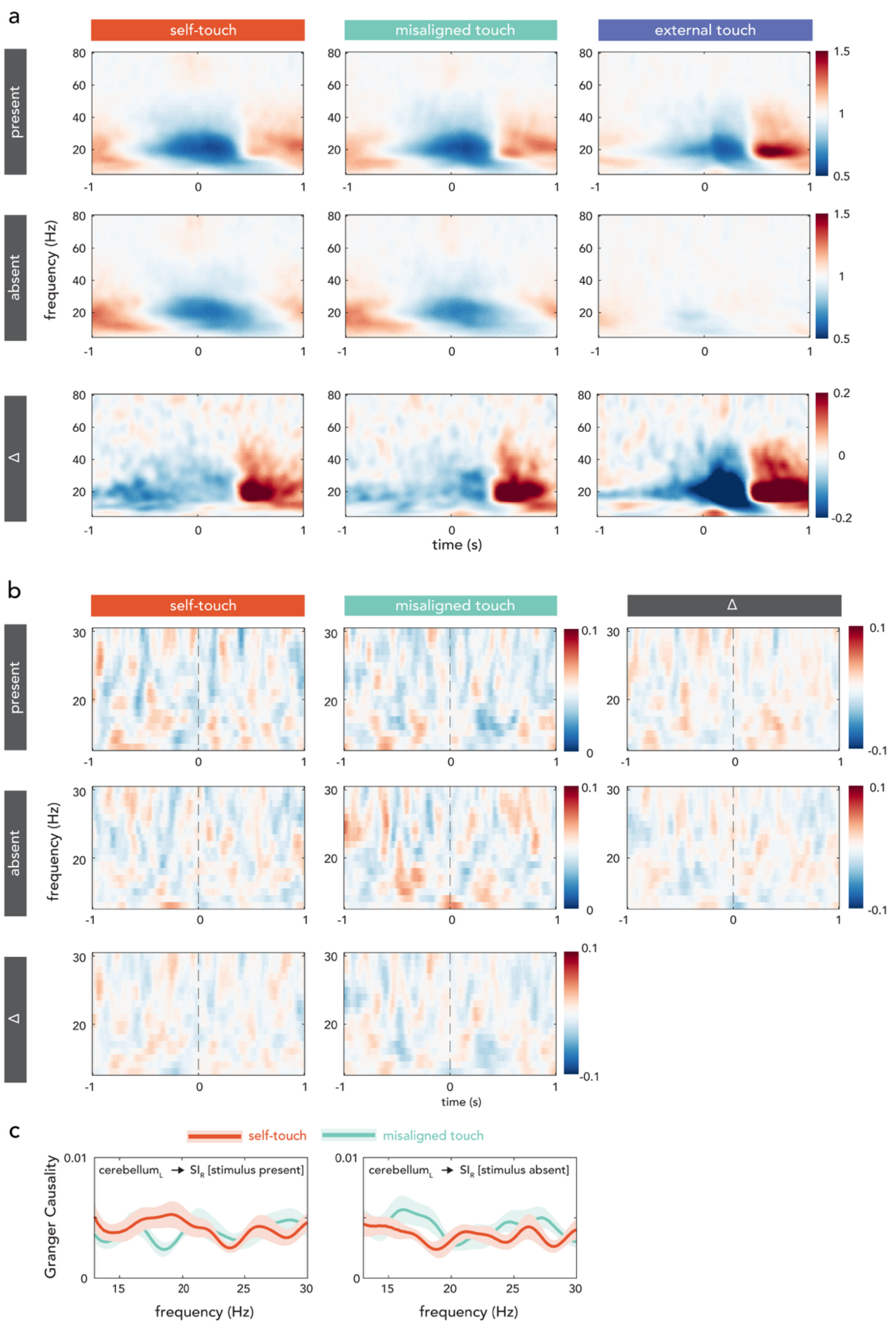


**Supplementary Figure S2. Time-frequency responses (TFR), coherence and directed connectivity. (a)** Grand-averaged TFRs at representative channels over the right sensorimotor areas for stimulus-present blocks (upper panel), stimulus-absent blocks (middle panel), and their difference (lower panel). **(b)** Grand-averaged coherence between virtual sensors in the left cerebellar lobule VI and the right primary somatosensory cortex in the beta band (13-30 Hz). Data are shown for the stimulus present blocks (upper panel) and stimulus absent blocks (middle panel), and their difference (lower panel: Δ = stimulus present – stimulus absent). **(c)** Non-parametric Granger causality values between virtual sensors in the left cerebellar lobule VI and the right primary somatosensory cortex (SI) in the beta band (13-30 Hz). Data are plotted separately for stimulus-present blocks (left panel) and stimulus-absent blocks (right panel). Shaded error bands represent ± standard error of the mean (s.e.m).

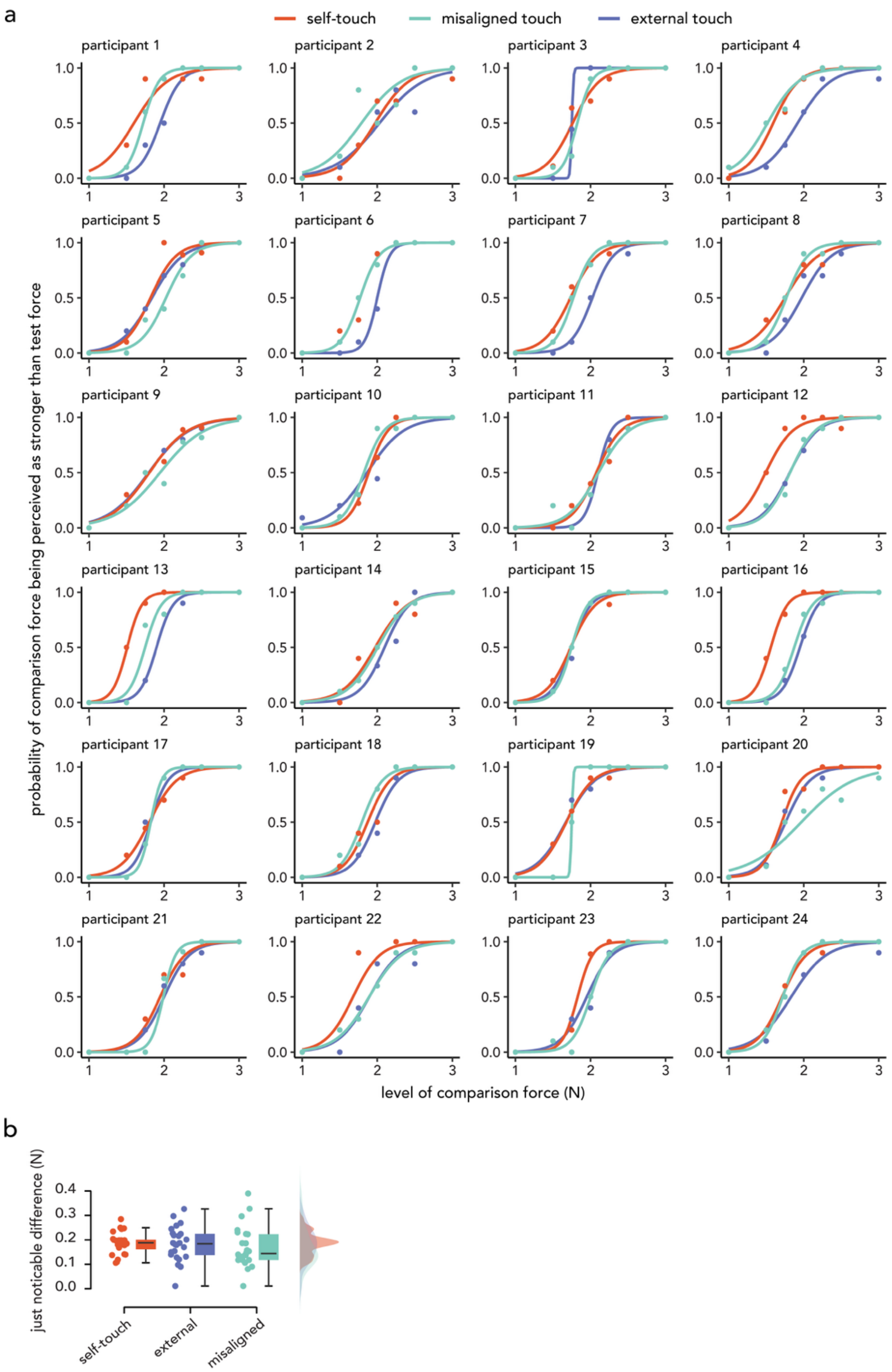


**Supplementary Figure S3. Force discrimination task participant psychometric fits and Just Noticeable Difference (JND) values**. **(a)** The fitted logistic models were very good, with McFadden's R-squared measures ranging between 0.600 to 0.970. **(b)** Comparable JNDs values in the three conditions of the behavioural session. The main effect of condition was not significant ($F(2, 46) = 0.57$, $p = 0.569$, $\eta_p^2 = 0.024$).
